# Metabolites reflect variability introduced by mesoscale eddies in the North Pacific Subtropical Gyre

**DOI:** 10.1101/2024.07.18.604132

**Authors:** William Kumler, Wei Qin, Rachel A. Lundeen, Benedetto Barone, Laura T. Carlson, Anitra E. Ingalls

## Abstract

Mesoscale eddies significantly alter open ocean environments such as those found in the subtropical gyres that cover a large fraction of the global ocean. Previous studies have explored eddy effects on biogeochemistry and microbial community composition but not on the molecular composition of particulate organic matter. This study reports the absolute concentration of 67 metabolites and relative abundances for 640 molecular features to understand how mesoscale eddies impact the metabolome of the North Pacific Subtropical Gyre during two cruises in 2017 and 2018. We find that many metabolites track biomass trends, but metabolites like isethionic acid, homarine, and trigonelline linked to eukaryotic phytoplankton were enriched at the deep chlorophyll maximum of the cyclonic features, while degradation products such as arsenobetaine were enriched in anticyclones. In every analysis, metabolites with the strongest responses were detected via untargeted mass spectrometry, indicating that the molecules most sensitive to environmental perturbation were not among the characterized metabolome. By analyzing depth variability (accounting for 20-40% of metabolomic variability across ∼150 meters) and the vertical displacement of isopycnal surfaces (explaining 10-20% of variability across a sea level anomaly range of 40 centimeters and a spatial distance of 300 kilometers), this analysis constrains the importance of mesoscale eddies in shaping the chemical composition of particulate matter in the largest biomes on the planet.

**Importance:** Mesoscale eddies are common ocean surface currents that circulate seawater vertically and horizontally. This stirring effect alters biogeochemistry and planktonic community composition. Here, we use metabolomics to determine how these eddy-induced changes influence the nature of organic carbon across an eddy dipole. We found that many small, polar molecules track with the overall particulate carbon in the system and that there were significant differences in metabolite composition between eddy states. A few metabolites reflected the increased importance of eukaryotic phytoplankton that were enriched by the higher nutrient supply from depth in the cyclonic eddies. Anticyclones contained more compounds that reflected a higher degree of degradation. This work answers outstanding questions about the importance of these common ocean features in shaping microbial community function.

## Introduction

High frequency observations at Station ALOHA in the North Pacific Subtropical Gyre (NPSG) over the past 25 years have revealed temporal and spatial variability in what had previously been considered a relatively homogenous environment (Karl 1999; Karl and Church 2017; Karl et al. 2021). A major source of variability comes in the form of mesoscale eddies in which water is entrained into circular surface currents tens to hundreds of kilometers in diameter (McGillicuddy 2016; Karl and Church 2017).

Eddies can be observed via satellite altimetry, which measures anomalies in sea surface height. Those that have a positive sea level anomaly (SLA) typically indicate regions where deep water layers and isopycnal surfaces are depressed and are associated with clockwise rotation in the Northern Hemisphere. In contrast, a negative SLA corresponds to counterclockwise rotation in the Northern Hemisphere and the uplift of deep water layers into the sunlit region of the ocean (McGillicuddy 2016). Mode-water eddies are an exception to the conventions established above (Sweeney et al. 2003; McGillicuddy et al. 2007) but here we only focus on the cyclonic (negative SLA) and anticyclonic (positive SLA) mesoscale features that are commonly observed near station ALOHA (Barone et al. 2019).

The uplift of deep, nutrient-rich seawater into the euphotic zone alters microbial communities (Rii et al. 2022). Measurements of chlorophyll in cyclones reveal a shallower and more intense maximum (Cornec et al. 2021; Barone et al. 2022) as a result of eukaryotic phytoplankton thriving in the higher nutrient concentrations while cyanobacterial biomass is reduced (Hawco et al. 2021). This growth of large eukaryotes corresponds to a net increase in biomass and primary productivity (Benitez-Nelson et al. 2007), especially near the deep chlorophyll maximum (Barone et al. 2022). In contrast, anticyclones produce conditions favorable to nitrogen fixation by accumulating diazotrophs such as *Crocosphaera* and *Trichodesmium* (Fong et al. 2008; Olson et al. 2015; Dugenne et al. 2020, 2023).

This shift in community structure and productivity should result in a corresponding shift in the composition of the particulate organic matter produced. Eukaryotic organisms have distinct chemical fingerprints from those of cyanobacteria and heterotrophic bacteria (Heal et al. 2021; Durham et al. 2022) and the relative contribution of these different taxa to total biomass should reflect their chemical composition. However, recent work from Harke et al. (2021) showed that overall community function may be robust across eddy types at the surface while Gleich et al. (2024) detected significant differences in protistan community composition and metabolic potential down to 250 meters, with heterotrophy-associated protistan transcripts enriched in the cyclone. The chemical composition of organic matter is one metric of community function since community metabolism and interactions are in part mediated through organic molecules, making their measurement useful for determining shifts in community dynamics.

In this work, we directly tested for differences in the chemical composition of particulate matter due to changes in eddy state in the NPSG. We sampled from cyclonic and anticyclonic eddies spatially near each other and used targeted and untargeted mass spectrometry to explore changes in the composition of small, polar molecules. We expected to find that overall metabolite abundance would reflect shifts in biomass over multiple depths. We also expected to see enrichment in the deep chlorophyll maximum of cyclonic features for those metabolites especially abundant in eukaryotic organisms. Finally, we predicted that these patterns would be robust across years and sampling regimes for a reliable way to link these ocean features with the chemical composition of organic matter in the open ocean.

## Results

We explored the impacts of isopycnal uplift and depression on the composition of particulate organic matter across pairs of mesoscale eddies of opposite polarity in the North Pacific Subtropical Gyre (NPSG). The first eddy pair was the focus of the July 2017 cruise, Microbial Ecology of the Surface Ocean-Simons Collaboration on Ocean Processes and Ecology (MESO-SCOPE, KM1709 aboard the R/V *Kilo Moana*). During the first phase of our two-phase expedition, a transect was taken from the north to the south consisting of 11 stations across a pair of eddies (the same sampling sites as in Barone et al. (2022) and Dugenne et al. (2023), Figure 1). Samples for particulate metabolomic analysis were collected from 15 meters, the deep chlorophyll maximum (DCM), and 175 meters at each station onto 0.2 µm filters. The next phase of the MESO-SCOPE cruise focused on Lagrangian sampling in the two eddy centers, starting with the cyclone (SLA = -14 cm) before progressing to the anticyclone (SLA = 24 cm). There, metabolomics samples were taken from the DCM as well as 10 and 20 meters above and below it. The second eddy pair was targeted during the March 2018 Hawaiian Eddy Experiment (HEE, FK180310 aboard the R/V *Falkor*). During this followup cruise, a strong anticyclone (SLA = 21 cm) and a nearby cyclone (SLA = -13 cm) were targeted for a similar set of biogeochemical measurements described in Dugenne et al. (2023) and Gleich et al. (2024) (Figure 1). Metabolomics samples were taken from the center of each eddy as described above at both 25 meters and the DCM. We analyzed these three different datasets separately and discuss the findings from each in sequence below.

**Figure 1:**
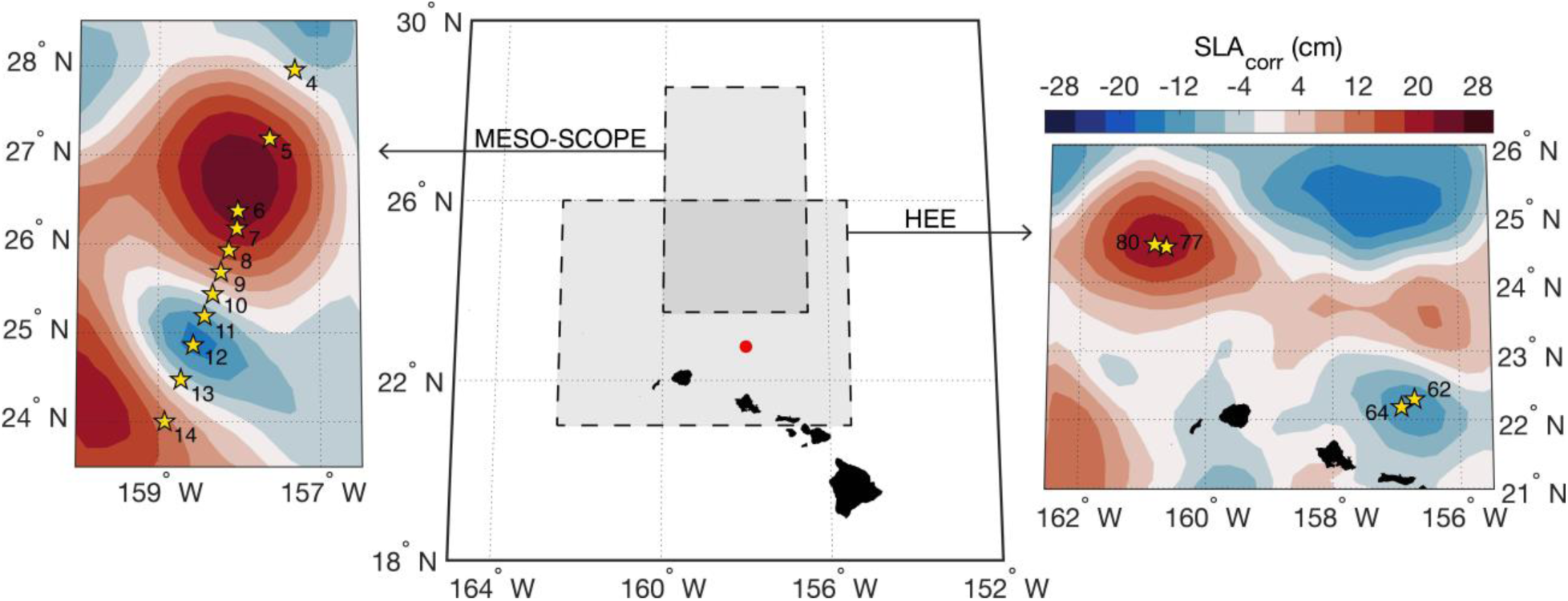
Sampling during MESO-SCOPE and the Hawaiian Eddy Experiment (HEE). Cruise bounds are shown in the large central map with Station ALOHA colored as a point in red near the Hawaiian Islands. Yellow stars denote sampling locations and station numbers relative to the sea level anomaly contours in the background during 28 June 2017 for the MESO-SCOPE cruise (left) and the 6 April 2018 for the HEE cruise (right). Lagrangian sampling near the eddy centers of the MESO-SCOPE cruise was performed by following drifters deployed in proximity of Stations 6 and 12.

### Metabolome variability across adjacent mesoscale eddies of opposing polarity during MESO-SCOPE explained by sea level anomaly variations

The metabolome clearly differed between cyclonic and anticyclonic samples, though the magnitude of this difference varied by depth. The largest differences in particulate matter composition were detected in the DCM and 175 meter samples, as shown by the NMDS in Figure 2. At the DCM, approximately 16% of the variation in the metabolome could be explained by sea level anomaly (SLA) alone (PERMANOVA, p = 0.003). Additionally, SLA was strongly correlated with the first principal component (PC) of the metabolome with a Pearson’s *r* of .615 and 27% of the variance explained by PC1 (Supplemental Figure 1A), indicating that this was one of the largest sources of variation in the dataset. Notably, the samples taken from the exact center of the cyclonic eddy at Station 12 were highly distinct and likely drove much of the explained variance. In the 175 meter samples, 12% of the variance was explained by SLA (PERMANOVA, p = 0.008) and SLA was highly correlated with the second PC of the dataset (r = .697, % variance explained by PC2 = 15.4, Supplemental Figure 1A), though the first PC did not seem to have any visible pattern with metadata and appeared to largely capture variation between biological triplicates (Supplemental Figure 1B). At 15 meters, SLA trends are less evident with a larger p-value (PERMANOVA, p=0.087) and a lower R^2^ (0.09) (Figure 2).

**Figure 2:**
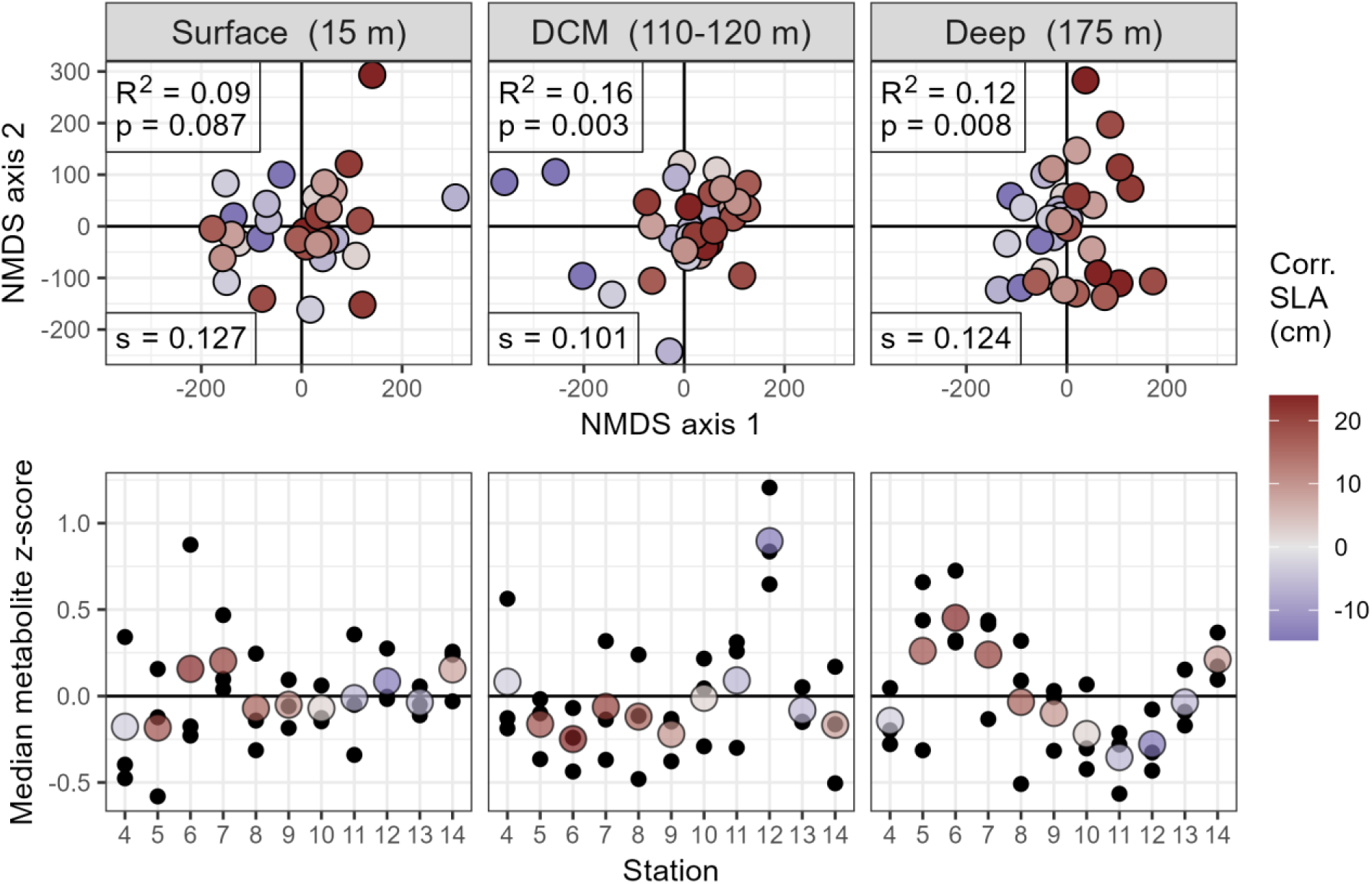
Distribution of metabolites in multivariate space across adjacent eddies of opposite polarity during the MESO-SCOPE transect, broken down by depth. Top panels depict non-metric multidimensional scaling (NMDS) plots with individual samples colored based on their corrected sea level anomaly. NMDS stress values (s) have been reported in the bottom left corner, while PERMANOVA R^2^ and p-values are reported in the top left. SLA trends are visible in the DCM and 175 meter samples, with dark blue circles consistently discriminating from the dark red circles along the first multidimensional axis. Bottom panels depict the direction and magnitude of this effect by plotting the mean value of all z-scored metabolites across three biological triplicates in color with the raw values in black behind.

**Figure 3:**
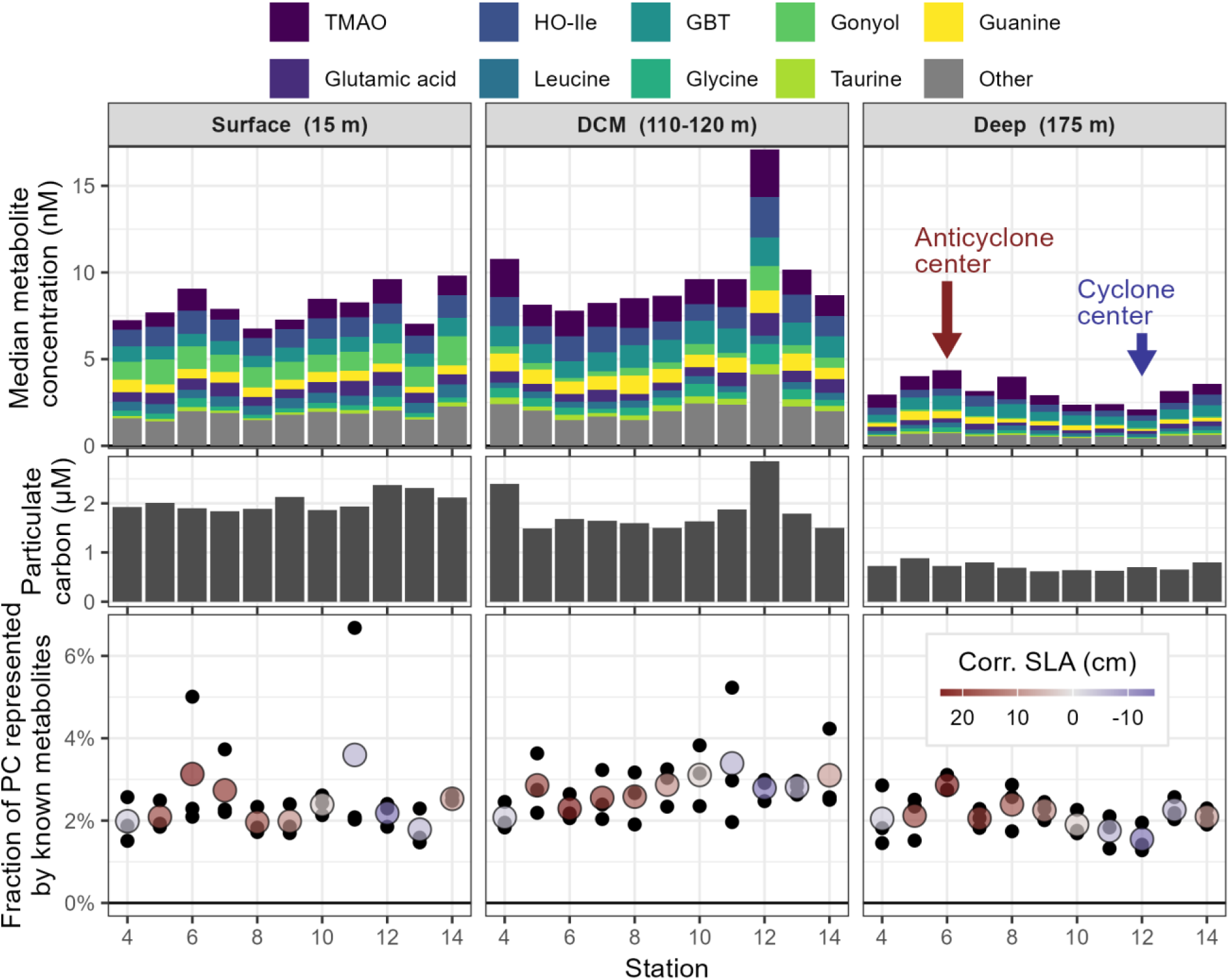
Differences in known metabolite concentration across the pair of adjacent eddies in the MESO-SCOPE transect separated by depth. Top panels depict concentrations of known compounds measured, where bar height corresponds to median triplicate concentration for each metabolite with the top 9 shown and the 44 other identified metabolites summed in grey. TMAO = trimethylamine N-oxide, HO-Ile = hydroxyisoleucine, GBT = glycine betaine, DCM = deep chlorophyll maximum. Center panels show the corresponding measurements of particulate carbon (PC), while the lower panels depict the fraction of total particulate carbon in the known metabolites. The mean value of three biological triplicates is shown in color and the raw values in black behind. Colors correspond to corrected sea level anomaly (Corr. SLA), with dark red indicating anticyclonic (positive SLA) and dark blue indicating cyclonic (negative SLA) eddy state.

To characterize the difference between eddies in a lower-dimensional space, we plotted an average metabolite peak area per station at each depth. We performed z-score normalization on each mass feature to give them all equal weight while preserving the variance between samples then calculated the median z-score for each sample, resulting in large positive values when metabolites are more abundant in a given sample than in the median sample (Figure 2). The average metabolite collected at the exact center of the cyclonic eddy (Station 12) had higher values than other samples at this same depth, indicating that most metabolites were more abundant at the cyclone’s DCM than elsewhere in the transect. The 175 meter samples, on the other hand, showed the largest metabolite abundances in the anticyclone while the cyclone had consistently lower median z-scores. This method highlighted the negative correlation between the average metabolite and SLA at the DCM (*r*_*DCM*_ = −.716) while the 15 meter and 175 meter samples showed the opposite trend (*r*_15*m*_ = .211, *r*_175*m*_ = .830).

This shift in metabolite abundance was largely driven by a corresponding shift in biomass. Particulate carbon and summed metabolite concentration were tightly correlated across all samples and depths (*β* = 24.9 ± 1.82 (SE), Pearson’s *r* = .813, p-value < 0.001, Supplemental Figure 2). This overall trend was largely driven by higher values at 15 meters and the DCM than at depth, but when analyzed at each depth individually the general correspondence held (Supplemental Figure 2). The DCM and 175 meter samples had the stronger trends (*β*_*DCM*_ = 22.6 ± 5.53, *r* = .593, p-value < 0.001; *β*_175*m*_ = 23.9 ± 7.36, *r* = .503, p-value = 0.003) while 15 meter samples showed no significant relationship (*β*_15*m*_ = 3.4 ± 12.9, *r* = .049, p-value = 0.792). This meant that metabolites contributed a relatively fixed fraction of carbon to the particulate pool with our 53 quantified metabolites representing between 1.5% and 5% of the carbon in the system (Figure 2). Particulate nitrogen was tightly correlated with particulate carbon (*r* = .933) but the fraction of particulate nitrogen represented by the quantified metabolites was slightly higher across the transect with contributions typically between 2% and 6%.

Although the untargeted pool contains more molecular features, we expect that a majority of the signal has been captured by our 53 targeted compounds. Of the total peak area that passed quality control, approximately 67% was captured by these 53 molecules. An additional 15% of the total peak area was putatively annotated as various inorganic ions for which no authentic standard was available but could be matched by mass and isotope pattern. Additionally, our targeted list accounted for 13 out of the top 20 largest molecular features by total peak area with an additional 4 features annotated as the inorganic ions for 17/20 of the largest features known. However, these calculations should be taken as estimates because peak area does not correspond directly to environmental concentration and it is entirely possible for abundant environmental compounds to have small peak areas if they ionize poorly on the mass spectrometer or are removed during the sampling and extraction process.

Samples taken from the DCM of the cyclone center (Station 12, Figure 2) showed especially high particulate carbon (67% increase, rising from 1.71 μM across all other DCM samples to 2.85 μM at Station 12) and the highest measured total metabolite concentrations (73% higher, increasing from 10.0 nM to 17.2 nM). This explains the large separation observed between these samples and the rest of the data in the NMDS plot of Figure 2.

The most abundant intracellular metabolite quantified in the eddy transect samples was trimethylamine N-oxide, with an average concentration of 1.38 nM and a distinct DCM maximum (Figure 2), though a few samples had concentrations an order of magnitude higher for unknown reasons (Supplemental Figure 3). A majority of the known molecules (37/53) had a similar pattern with a subsurface maximum at the DCM, including glycine betaine, glutamate, and guanine. Thirteen molecules obeyed a different pattern and decreased monotonically with depth, including the molecules hydroxyisoleucine, gonyol, and dimethylsulfoniopropionate. Two known molecules (arsenobetaine and O-acetylcarnitine) increased with depth, potentially representing particle degradation during export.

### Particulate organic matter compositional shifts across eddy transect

In addition to absolute shifts in metabolite concentration due to biomass differences, we also explored compositional shifts in the metabolomes by assessing the fraction of the total metabolite pool contributed by each compound across the eddy transect. This allowed us to isolate the signal due to shifts in community composition and organismal response from those introduced by changes in biomass of the community as a whole.

In the 15 meter surface samples, only seven compounds had significant changes in relative peak area across the eddy transect, two of which were known: 5-oxoproline, which decreased from 0.2-0.3% of the total peak area at the center of the anticyclone to 0.05-0.1% in the cyclone; and a combined peak of sarcosine and beta-alanine which increased from 0.01% to 0.02-0.03%. The largest shift significant at an α = 0.05 level was an unknown mass feature with *m/z* = 189.12338 and a retention time around 8.6 minutes, the relative contribution of which increased from approximately 0.2-0.3% in the cyclone to ∼1.5% in the anticyclone.

At the DCM, twenty-two compounds changed significantly in relative peak area across the transect. Seven of these were known metabolites, of which trigonelline and homarine were most enriched in the cyclone while arsenobetaine was the only metabolite with a larger fraction of the total peak area in the anticyclone. Homarine showed a very large and highly significant shift, representing about 2.5% of the total peak area in the anticyclone but 7-8% in the cyclone center. Trigonelline (N-methyl niacin), an isomer structurally very similar to homarine but biologically distinct, had a surprisingly strong correlation with homarine (Pearson’s *r* = .855). Its peak areas were consistently around one third of homarine’s but still large enough to have the second-largest shift in relative peak area of the significantly different compounds. Similarly, arsenobetaine represented about 0.1% of the peak area in the cyclone and 0.5-0.6% of the total peak area in the anticyclone. The most significantly different compounds at the DCM in each direction, however, were both unknowns. The lowest p-value (6.4 × 10^−6^ after FDR correction) in the DCM data was a mass feature with an *m/z* of 173.09211, a retention time also around 8.6 minutes, and enrichment in the anticyclone with a putative chemical formula of [M+H] = C_7_H_13_N_2_O_3_, possibly glycylproline or prolylglycine. The next-lowest (p = 2.5 × 10^−4^ after FDR correction) was enriched in the cyclone and had an *m/z* of 275.0712 and a retention time of 11.4 minutes with a putative formula of C_18_H_11_O_3_.

Finally, in the 175 meter samples we detected 43 mass features with a significant association with SLA. Notably, all detected nucleobases (guanine, adenine, and cytosine) were positively associated with SLA (higher values in the anticyclone than the cyclone), though the strongest associations were found in O-acetylcarnitine, betonicine, and tyrosine. Both O-acetylcarnitine and betonicine effectively quintupled their contribution to total peak area, shifting from approximately 0.1% to ∼0.5% over the transect. Acetylcholine was the only known compound more relatively abundant in the cyclone than in the anticyclone along with one unknown of *m/z* 131.0340. The unknown at *m/z* 173.0921 with the strongest trend at the DCM again had the strongest trend in the 175 meter samples, here with an FDR-corrected p-value of 2.0 × 10^−7^.

### High vertical resolution sampling near eddy center DCM reveals distinct metabolome between cyclone and anticyclone

We further investigated the metabolomic response to SLA at the DCM by collecting samples taken at high-resolution depth intervals around the DCM at the center of both eddy poles. Here, we found that this ∼40 meter depth range and the transition between the eddies explained similar amounts of variation in the data (R^2^_*depth*_ = 0.253, R^2^_*SLA*_ = 0.238) and were both highly significant factors (permutational p-values << 0.001, Figure 4). Additionally, both sample depth and SLA were correlated with the first principal component of the metabolite matrix (*r*__*depth*__ = .766, *r*__*SLA*__ = .684, fraction of variance explained by PC1 = 30.8%). The cyclonic samples in particular show a much larger spread than the anticyclonic ones, indicating larger sample variability in the cyclone DCM relative to the anticyclone. A depth gradient is visible along NMDS 2, with the deepest samples generally at the top right of the plot and the shallowest ones closer to the bottom (Figure 4).

**Figure 4:**
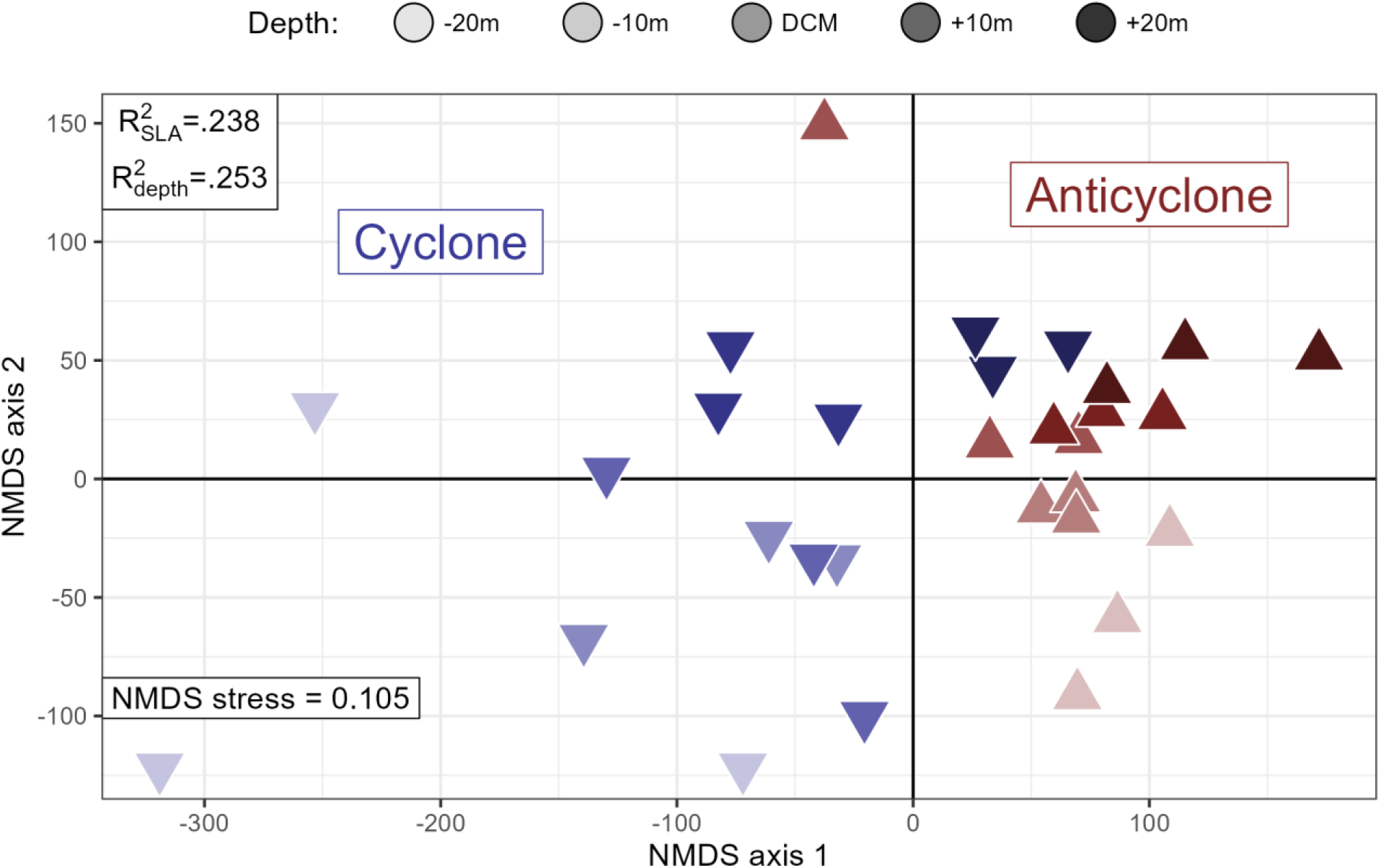
NMDS plot of high-resolution depth sampling around the deep chlorophyll maximum (DCM, ∼115 meters) at the two eddy centers during MESO-SCOPE. Red upward-pointed triangles are from the anticyclone and blue downward-pointed ones are from the cyclone. Shading intensity reflects the depth above or below the DCM. PERMANOVA estimates of the variance explained by depth and sea level anomaly (SLA) are noted in the upper left corner and the NMDS stress value is reported in the bottom left.

To characterize the observed differences in multivariate space we used k-means clustering as an unsupervised way to identify dominant trends within the data (Figure 5). We found that a majority of the metabolites (Clusters 1 and 2, 63% of the total) fell into clusters with larger peak areas in the cyclone, as expected, while also detecting 32 metabolites that clustered such that the mean metabolite was enriched in the anticyclone (Cluster 4, Figure 5). Cluster 2 had a distinct decrease in relative peak area with depth, while the other clusters had much less clear depth trends. Cluster 1 also showed a DCM maximum for the samples from the anticyclone, correlating well with flow cytometry counts of picoeukaryotes (Pearson’s *r* = .851) that was not present in the cyclone (*r* = .013).

**Figure 5:**
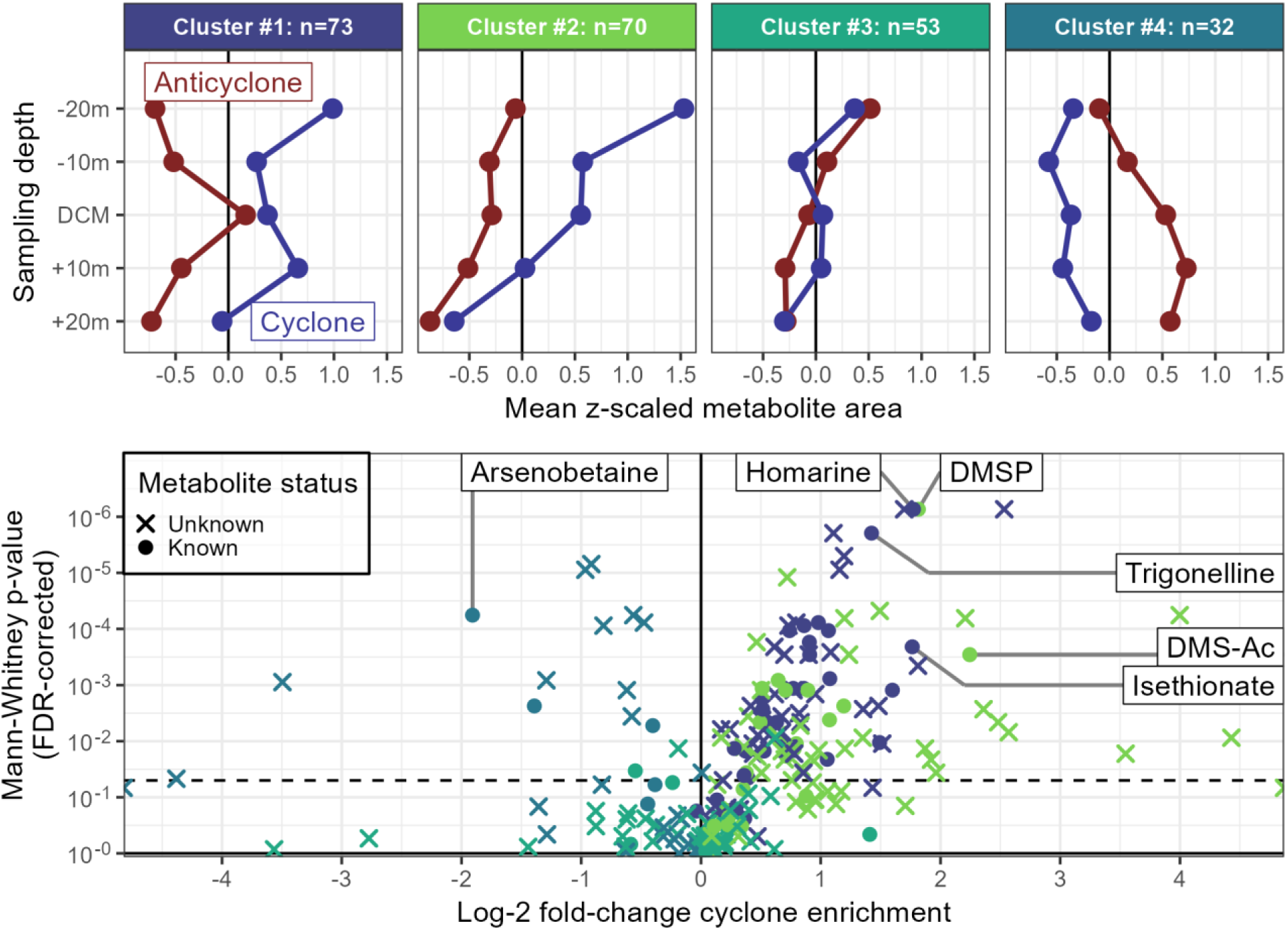
Distribution of metabolites in the high-resolution depth samples from the centers of each MESO-SCOPE eddy. The upper row of plots shows k-means clusters where points denote the average z-scored peak area for both known and unknown metabolites across the samples and are colored by the eddy from which they were taken. Clusters have been ordered by number of metabolites in each group and the total is denoted in the panel titles. Both depth trends (mostly a net decrease in metabolites with depth) and eddy effects (cyclonic enrichment in clusters 1 and 2, anticyclone enrichment in cluster 4) are observable. The lower plot shows the individual known and unknown metabolites where points correspond to the FDR-corrected p-value estimated by the nonparametric Mann-Whitney U test and the log_2_ fold-change calculated with the average peak area in the cyclone divided by the average peak area in the anticyclone. Colors have been assigned using the k-means clusters and shapes have been assigned based on the status of the mass feature as either a known metabolite that was matched to an authentic standard or an unidentified metabolite. The dashed line across the figure represents the 0.05 level of significance as a visual cue for metabolites above which the differences between the eddies are unlikely to be due to chance. DMSP = dimethylsulfoniopropionate, DMS-Ac = dimethylsulfonioacetate

The majority of the individual metabolites in these high-resolution depth profiles differed significantly between the two eddies (121/228, α = 0.05). Of those, 15 were enriched in the anticyclone and 106 were more abundant in the cyclone. As expected, all compounds enriched in the anticyclone were part of Cluster 4 and all those enriched in the cyclone belonged to either Cluster 1 or 2 (Figure 5).

Many of the known metabolites enriched in the cyclone function as osmolytes in the cell and might be enriched due to the increased eukaryotic phytoplankton biomass. However, some metabolites such as isethionate, the reduced sulfur osmolytes dimethylsulfoniopropionate (DMSP) and dimethylsulfonioacetate (DMS-Ac), and the isomers homarine and trigonelline, were enriched in excess of biomass (Figure 5). A few metabolites more abundant in the anticyclone were also given putative identifications based on RT and *m/z* matching with internal standards run at a later time on the mass spectrometer. Of these, the putative arsenobetaine was the most significantly different among these with a peak area at the anticyclone DCM nearly quadruple that of the cyclone (Figure 5). Of note, the 173.0921 *m/z* mass feature noted above as strongly enriched in the anticyclone was among the most significantly different between the two eddies, along with two isomers at an *m/z* of 170.1176 and retention times around 8-9 minutes that increased by a factor of 1.5 in the anticyclone (putative formula C_7_H_14_N_4_O).

### Hawaiian Eddy Experiment data reveals a different community and response to mesoscale eddies

Given the strong signals detected in the samples from the MESO-SCOPE cruise both across the eddy transect as well as in the high-resolution DCM sampling, we expected to find similar results in an analogous dataset. Samples were collected during a 2018 cruise on the R/V *Falkor* that again targeted both a cyclonic and an anticyclonic eddy in the North Pacific Subtropical Gyre near Station ALOHA as part of the Hawaiian Eddy Experiment (HEE).

The HEE data was characterized by high inter-replicate variability relative to the samples from the MESO-SCOPE cruise, with 25 meter samples in particular highly variable in multivariate space (Figure 6). Despite this, we still saw the importance of sea level anomaly as a significant explanatory factor in the dataset (R^2^*_SLA_* = 0.090, p-value = 0.017) as well as the larger depth differences between the surface (25 meters) and DCM (110 - 120 meters) (R^2^*_depth_* = 0.251, p-value << 0.001).

**Figure 6:**
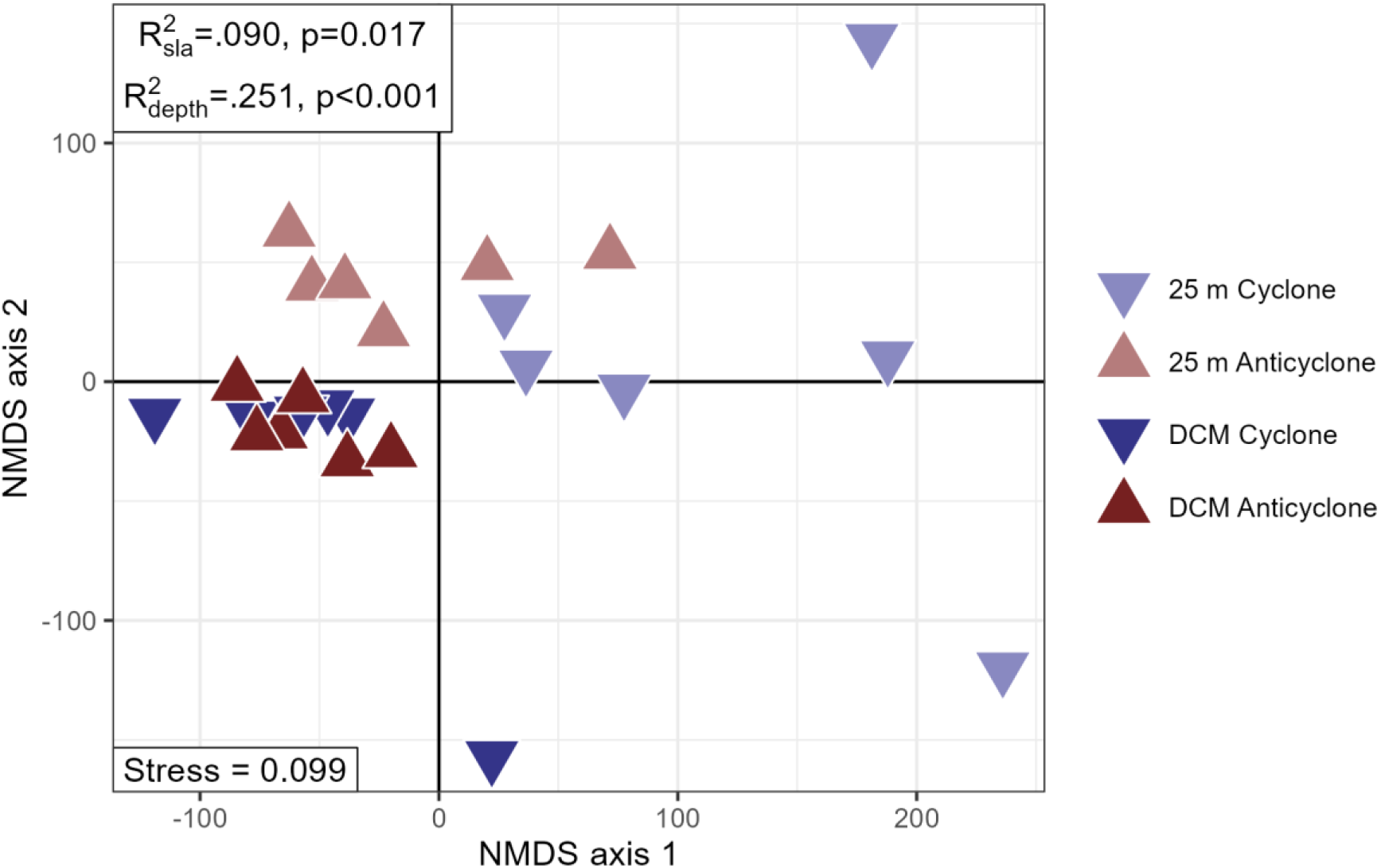
Non-metric multidimensional scaling (NMDS) plot from the Hawaiian Eddy Experiment cruise data, in which points correspond to individual samples and have been colored and shaped by their source eddy status and shaded by the depth from which they were collected. The NMDS stress value has been reported in the bottom left corner, while PERMANOVA R^2^ and p-values are reported in the top left. Samples from 25 meters deep are visibly distinct from the deep chlorophyll maximum (DCM) samples and an SLA signal is visible in the 25 meter samples only.

When analyzed separately as distinct depths, the 25 meter samples had higher variances explained by eddy (R^2^=.26) and a lower p-value (0.013), while the DCM samples were no longer likely to be distinct between the two eddies (p-value = 0.171) (Figure 6, Supplemental Figure 4).

We were also able to use the HEE dataset to test whether the compounds detected as significantly different in the MESO-SCOPE dataset were also different in this eddy pair. We expected to find abundant TMAO and more hydroxyisoleucine at the surface than the DCM in addition to most compounds enriched in the cyclone DCM with biomass, especially the six compounds mentioned above in the results of Figure 5 (trigonelline, homarine, DMS-Ac, DMSP, taurine, and isethionic acid).

As expected, TMAO was again one of the most abundant metabolites detected in the particulate matter with concentrations around 1.2 nM at 25 meters and 0.4 nM at the DCM, second only to the high levels of the amino acid glycine (2 nM at 25 meters, 0.9 nM DCM), though the differences between 25 meters and DCM were not significant for either compound. We also detected a significant difference between depths for hydroxyisoleucine (25 meter mean = 0.32 nM, DCM mean = 0.21 nM, t-test p-value=0.022 with n=12 samples at each depth). These results imply that the overall depth structure of the metabolome of the gyre is relatively fixed for the most abundant compounds.

The eddy effects, on the other hand, were much less strong during this cruise. Of the six known metabolites with the strong enrichment in the cyclone at the DCM in the MESO- SCOPE cruise, only isethionic acid was significantly different in the HEE dataset (isethionic acid t-test p-value_*FDR*_ = 0.043, other five p-values > 0.25). This difference in isethionic acid concentration was only found after normalizing each sample to the sum of all metabolites in the sample to control for biomass. Arsenobetaine was again found to be strongly enriched in the anticyclone at the DCM both when normalizing to biomass and when not, just as in MESO-SCOPE. Surprisingly, the 173.0921 *m/z* mass feature was also slightly enriched but this time in the *cyclone* DCM with peak areas approximately 1.4 times the values in the anticyclone (t-test p-value = 0.067, Mann-Whitney p-value = 0.045). The 170.1176 *m/z* mass feature was not detected in the HEE samples.

## Discussion

### Multivariate approaches reveal metabolome-wide shifts across pairs of eddies of opposite polarity

We measured the metabolome of samples collected across two sets of adjacent eddies of opposite polarity to explore the effect of sea level anomaly (SLA) on metabolite composition and found that the altered biogeochemistry and microbial community composition between cyclonic and anticyclonic eddies explain a significant portion of the observed variations in their particulate metabolomes.

In all of the datasets analyzed here, we detected a significant difference in the composition of the metabolome between the adjacent eddies. This effect was strongest in the samples taken during the Lagrangian stations at the center of each eddy in the 2017 MESO-SCOPE cruise, with nearly a quarter of the total variance explained by the eddy from which the samples were taken. In the samples taken along the eddy transect, the largest effect was detected in the deepest samples taken from 175 meters and from the DCM, with much less of a response at 25 meters. In the HEE samples, however, the opposite was true with a larger SLA response at 25 meters than that at the DCM. In each case, the differences introduced by SLA were smaller than those of depth when compared directly, with even the high-resolution sampling around the DCM finding slightly more variance explained by the 40 meter difference in sampling depth than the 40 centimeter difference in SLA. This result agrees well with prior research by Heal et al. (2021) and Kumler et al. (2023) showing the large effect of sampling depth on the metabolome.

Although possible that the observed differences in metabolomes across adjacent eddies differing in polarity are due to latitudinal shifts or simply background variation in the gyre environment, the transect data in particular implies that this is unlikely to be the case. Samples taken when the absolute value of the SLA was less than 5 cm were more similar to each other than to those samples taken within the eddies despite larger differences in latitude. The transect sampled both outside of each eddy and between the two across a large spatial gradient (∼4° latitude and 2° longitude), yet the non-eddy (absolute value of SLA < 5 cm) samples from all three locations grouped together, showed similar median metabolite concentrations, and were visually similar in the most abundant metabolite composition. These results imply that strong cyclonic and anticyclonic eddies represent endpoints in NPSG composition, in agreement with results from Barone et al. (2019) which found the most extreme values in the Hawaii Ocean Time series data typically detected when eddies passed over Station ALOHA. We also detected the largest differences in the DCM metabolome at the exact center of the eddy, with Station 12 highly distinct from even the nearest stations, while no such stark difference was found in the anticyclone (Figures 1 and 2). This perhaps indicates that the exact center of the cyclone has a unique metabolome at the DCM relative to the rest of the transect, rather than existing as a smooth continuation.

Shifts in total biomass were a large driver of metabolomic differences, with earlier work in the same location by Barone et al. (2022) showing that particulate carbon, chlorophyll, and beam attenuation were all 20-80% higher in the DCM of the cyclone relative to the anticyclone. We see this reflected particularly well in the targeted metabolites measured here, with clear depth trends and a strong correlation between total metabolite concentration and particulate carbon values. However, even after controlling for biomass effects by normalizing each sample to the sum of signal measured within it we still found a significant SLA effect, likely due to shifts in the community composition and in particular the increased eukaryote presence in the cyclone DCM.

The NMDS plots of the high-resolution depth samples were also illustrative of sample dissimilarity with clear SLA and depth trends. The anticyclonic samples grouped tightly and showed little variance with depth or triplicate when compared to the cyclonic samples with the exception of DCM triplicate B, which was highly distinct for unknown reasons. In contrast, the cyclonic samples were much more variable as is perhaps expected when the biomass is in the form of larger phytoplankton that are less homogenous in the environment. One notable aspect of their clustering was the way in which the deepest samples (DCM plus 20m) grouped most closely to the anticyclonic samples, perhaps indicating that below the DCM the metabolome rapidly approaches a uniform deep-water signal. This result was noted previously in the 175 meter samples of the eddy transect in Kumler et al. (2023), where samples from the deep euphotic zone showed greater intra- depth similarity than samples from the DCM or 25 meters.

Given the confluence of both depth and SLA signals and the way that the boxplots of Figure 2 would confound the SLA signal with the depth variance, we used k-means clustering to group similar compounds and provide a reduced dimensionality space for visualization.

This revealed four major patterns of metabolite response, with a majority of compounds responding to eddy state (Clusters 1, 2, and 4 in Figure 5). It is interesting to note that the sole cluster in which abundances in the anticyclone are greater than those of the cyclone is also the only cluster to generally increase in abundance with depth (Cluster 4), perhaps indicating that compounds more concentrated in the anticyclone are the same kind of degradation products and recalcitrant carbon typically found at depth.

The Hawaiian Eddy Experiment (HEE) data upset many of the expectations we developed during the MESO-SCOPE cruise in the previous year. Most surprising was the large difference between the eddies detected at the *surface*, while the DCM samples were functionally indistinguishable. This contrasts directly with the results from MESO-SCOPE and the results in Gleich et al. (2024), who found eddy-driven shifts in protistan community composition to be larger at depth than at the surface. However, Dugenne et al. (2023) found differences in nitrogen fixation rate and nitrogen fixer composition varying widely throughout the upper water column. The large surface differences were especially surprising given the large inter-replicate differences between the HEE surface samples in this study, while the DCM samples tended to be much more consistent.

### Univariate approaches highlight individual metabolites responding to lifted and depressed isopycnals

Several molecule- or pathway-specific narratives emerged from the metabolomic data. The untargeted approach used here allowed us to describe and characterize signals from small molecules whose identity was unknown, a particularly promising approach in open ocean gyres where the largest fraction of unknowns is found (Heal et al. 2021). We found that our list of authentic standards covered fewer metabolites in anticyclonic eddies at all depths, with the molecular diversity of the cyclone much better characterized. Many of the trends detected persisted even after normalizing within each sample, indicating that the shifts discussed have implications beyond simple scaling with biomass.

Of those molecules whose identity was known, the clearest response to SLA was that of the enigmatic osmolyte homarine at the DCM. This abundant compound increased approximately threefold in concentration from < 50 pM in the anticyclone to around 150 pM in the center of the cyclone in both the eddy transect and eddy center datasets. The pattern was weaker in the noisier HEE data but the largest concentrations were still detected in the cyclone and the median cyclone measurement was greater than the maximum value for the anticyclone. This molecule has a well-established role as an osmolyte in eukaryotic phytoplankton and cyanobacteria (Gebser and Pohnert 2013; Dawson et al. 2020; Heal et al. 2021; Durham et al. 2022) as well as a documented decrease in abundance with depth (Heal et al. 2021), indicating that its response to SLA is potentially due to differences in physical and chemical attributes that in turn control biomass, community composition, and recycling rates. Curiously, the distribution of homarine was also tightly correlated with that of trigonelline. Although they have structural similarity they are not known to have a relationship beyond their shared function as osmolytes.

Isethionate is a known osmolyte thus far found exclusively in eukaryotes (Durham et al. 2022) and strongly associated with a few diatoms in particular (Heal et al. 2021). This compound was enriched in the cyclone of the DCM in both the MESO-SCOPE and HEE cruises along with its precursor taurine, validating earlier results from Barone et al. (2022) that showed strong enrichment of eukaryotic phytoplankton at the DCM and *Pseudo-nitzschia* in particular.

The strong SLA signal detected among the 175 meter samples was partially surprising to us given our expectation about the strongest effect at the DCM where eddy effects lift large concentrations of nutrients above the 1% light level. Instead, we found more compounds overall to be significantly different across the eddy transect in the 175 meter samples than we did at the DCM (α=0.05, 43 compounds at 175 meters vs 22 at the DCM). This was likely driven by the enhancement of organic matter and heterotrophic picoplankton below the DCM as seen in Barone et al. (2019), with the enrichment of nucleobases in the 175 meter samples additionally hint at increases in bacterial biomass given that nucleobases tend to be highly abundant in bacterial cultures (Heal et al. 2021). The abundance of a mass feature putatively identified as acetylcholine in the 175 meter cyclone samples followed the same general trend as arsenobetaine, both enriched at depth and in the anticyclone, and may be another compound that results from heterotrophic degradation of phytoplankton metabolites (Durham et al. 2022). These results raise important questions about the depth at which the community is isolated from SLA effects, with additional data from Barone et al. (2022) and Gleich et al. (2024) indicating that even at 250 meters eddy effects are still discernable.

Differences between the two cruises could be due to differences in eddy age and/or the different period of the year when sampling took place during the two cruises (March/April in 2018 instead of the June/July sampling in 2017) resulting in shifts in community composition and response. Barone et al. (2022) documents the 2017 cruise community composition, reporting abundant Prymnesiophyceae at both 15 meters and the DCM, with other significant contributions from dinoflagellates, diatoms, and green algae. In the 2018 cruise both haptophytes and dinoflagellates again dominated the eukaryotic fraction, though surprisingly little was contributed by diatoms (Gleich et al. 2024). Dugenne et al. (2023) and Harke et al. (2021) also highlight uncontrolled differences between the two sets of eddies based on their origin and age.

Missing links between the environment with its associated community and the composition of particulate matter make it difficult to estimate this important control on the marine carbon cycle. This is especially true for dynamic and short-lived environments such as eddies. Metabolites offer a way to bridge this gap in knowledge but the relative impact of eddies on their contribution and fate was not previously estimated. In this work, we have shown the sensitivity of the metabolome to various environmental and community factors and contrasted it to the well-established effect of depth to constrain the importance of including eddy effects in models of elemental cycling.

## Conclusions

This study reports how changes in the biogeochemistry and community composition due to mesoscale eddies affect the metabolome and thereby the composition of particulate organic matter. A transect across adjacent mesoscale eddies of opposite polarity showed that many metabolites track closely with metrics of overall biomass, with eddy effects stronger at 175 meters than at 15 meters or the deep chlorophyll maximum (DCM). High-resolution depth sampling of the DCM at the center of each eddy elucidated several known and unknown biomarkers likely corresponding to the previously documented increase in eukaryotic phytoplankton in the cyclone. Metabolite clusters aligned well with expected trends in abundance with depth and between the two eddies, with convergence below the DCM towards a general deep-sea metabolite signal for many metabolites. By contrasting these effects with a follow-up analysis in the centers of a separate pair of mesoscale eddies of opposite polarity the following year, we learned that the impact of eddies may be influenced by their origin, age, and period of the year. Finally, the metabolites with the strongest responses to eddy effects were unknown, and a relatively smaller proportion of known metabolites in the anticyclone demonstrates the utility of untargeted metabolomics in exploring environmental variation and the large amount of information that is missed by only investigating known compounds.

## Acknowledgements

The authors would like to thank the other members of the Ingalls Lab who provided assistance in sample processing, integration, and analysis. We are also grateful to the SCOPE science team (Tara Clemente and Tim Burrell) for sample collection during the Hawaiian Eddy Experiment. We would like to acknowledge the captain and crew of the R/V *Kilo Moana* and R/V *Falkor* during the KM1709 and FK180310 cruises for making this science possible. We would also like to acknowledge the analysts involved in the biogeochemical measurements for their willingness to provide data and answer questions (Eric Grabowski, Karin Bjorkman, and Rhea Foreman). This work was supported by grants from the Simons Foundation (SCOPE Award ID 329108 to AI, SF Award ID 385428 to AI, Postdoctoral Fellowship in Marine Microbial Ecology ID 548565 to WQ)

The authors have no conflicts of interest to declare.

## Data availability

Data and code used for this manuscript are all available online. Biogeochemical data can be sourced from the Simons SCOPE website at https://scope.soest.hawaii.edu/data/ and metabolomics data has been uploaded to Metabolomics Workbench under Project ID PR001738, where the data can be accessed directly via its DOI: 10.21228/M82719. Scripts used to process metabolomics data are available on GitHub at https://github.com/wkumler/MesoscopeMetabolomicsManuscript.

## Methods

### Cruise information

Samples were collected from two cruises in the North Pacific Subtropical Gyre near Station ALOHA that targeted strong mesoscale eddy features as described in Dugenne et al. (2023) and Gleich et al. (2024). Briefly, the 2017 MESO-SCOPE cruise (Microbial Ecology of the Surface Ocean-Simons Collaboration on Ocean Processes and Ecology, KM1709 on the R/V *Kilo Moana*) consisted of a transect across adjacent cyclonic and anticyclonic eddies as well as two long-term Lagrangian stations at the center of each eddy. The cyclonic eddy had a maximum negative sea level anomaly (SLA) of -20 centimeters and the anticyclonic eddy reached +24 centimeters. The transect samples were taken at various times of day as the ship transected the adjacent eddies while the eddy center samples were all collected between 5 and 8 pm. In 2018, the Hawaiian Eddy Experiment (HEE, FK180310 on the R/V *Falkor*) targeted new cyclonic and anticyclonic eddies in approximately the same location with samples taken at the center of each eddy (maximum negative anomaly in the cyclone = -15 cm, maximum positive anomaly in the anticyclone = +26 cm) (Figure 1).

### Biogeochemical data

Biogeochemical measurements were collected as described in Barone et al. (2022) and Dugenne et al. (2023) mostly following protocols used by the HOT program (http://hahana.soest.hawaii.edu/hot/methods/results.html). Briefly, all environmental data was collected via CTD rosette except SLA which was measured via satellite. Particulate carbon and particulate nitrogen were measured using an elemental analyzer. Nitrate + nitrite (N+N) was measured on an autoanalyzer except where concentrations were below 100 nM in which case they were measured using chemiluminescence. Soluble reactive phosphorus (phosphate, PO^3−^) was measured on an autoanalyzer or following the magnesium induced coprecipitation method. Values were interpolated linearly to the depths at which metabolite samples were taken using all data collected at the given station. Two particulate carbon values were determined to be spurious, with concentrations 2-3 times higher than typical Station ALOHA values, and were instead estimated using a best-fit linear regression against beam attenuation.

### Sample collection for particulate metabolites

Samples were obtained using the onboard CTD rosette to collect water from 15 meters, the deep chlorophyll maximum (DCM), and 175 meters during the MESO-SCOPE eddy transect and from the DCM ±10 and ±20 meters during the eddy center sampling in the MESO- SCOPE cruise and from 25 meters and the DCM during the HEE cruise. The DCM was determined visually from fluorometer data during the CTD downcast and Niskin bottles were tripped during the return trip to the surface. Seawater from each depth was sampled in triplicate by firing one Niskin bottle for each sample. Samples were brought to the surface and decanted into prewashed (3x with DI, 3x with sampled seawater) polycarbonate bottles for filtration. 10L samples were filtered by peristaltic pump onto 142mm 0.2 µm Durapore filters held by polycarbonate filter holders on a Masterflex tubing line. Pressures were kept as low as possible while still producing a reasonable rate of flow through the filter, approximately 250-500 mL per minute. Samples were then removed from the filter holder using solvent-washed tweezers and placed into pre-combusted aluminum foil packets that were then flash-frozen in liquid nitrogen before being stored at -80 °C until extraction. A methodological blank was also collected by running filtrate through a new filter and then treated identically to the samples.

### Metabolite sample extraction

Extraction followed a modified Bligh & Dyer approach as detailed in Boysen et al. (2018). Briefly, filters were randomized and added to PTFE centrifuge tubes with a 1:1 mix of 100 µm and 400 µm silica beads, approximately 2 mL -20 °C Optima-grade dichloromethane, and approximately 3 mL -20 °C 1:1 methanol/water solution (both also Optima-grade). Isotope labeled extraction standards were also added during this step (Supplemental Table 1). The samples were then bead-beaten three times, followed by triplicate extraction of the aqueous layer and addition with fresh methanol/water mixture with additional bead- beatings in between. Samples were then dried down under ultrapure nitrogen gas and warmed using a Fisher-Scientific Reacti-Therm module. Dried aqueous fractions were reconstituted in 380 µL Optima-grade water and amended with 20 µL isotope-labeled injection standards (Supplemental Table 1) to measure the variability introduced by chromatography and ionization. The reconstituted fraction was syringe-filtered into precombusted glass inserts in HPLC vials to remove any potential clogging material. Samples were then additionally diluted 1:1 with Optima-grade water to prevent overloading on the column and reduce salt effects.

A pooled sample was created by combining 20 µL of each sample into the same HPLC vial, and a 1:1 dilution with water created a half-strength pooled sample from that to assess matrix effects and obscuring variation (Boysen et al. 2018). Also run alongside the environmental samples were samples of authentic standards split into two mixes (Supplemental Table 2). Standards were dissolved in Optima-grade water to help with identification and into an aliquot of the pooled sample to quantify matrix response factors. HPLC vials containing the samples were frozen at -80 °C until thawing shortly before injection.

### HPLC-MS methods

Separate HPLC-MS runs were used for the MESO-SCOPE eddy transect, the MESO-SCOPE eddy center, and the HEE cruise data. Eddy center samples were run in February 2018, eddy transect samples were run later during August of that year, and the HEE samples were run in July 2019. Each run was treated as a single batch and injected in sequence while maintaining the randomization that had occurred during extraction to minimize chromatographic shifts from solvent or column switching.

For each run, a Waters Acquity I-Class UPLC with a SeQuant ZIC-pHILIC column (5 µm particle size, 2.1 mm x 150 mm, from Millipore) was used with 10 mM ammonium carbonate in 85:15 acetonitrile to water (Solvent A) and 10 mM ammonium carbonate in 85:15 water to acetonitrile (Solvent B) at a flow rate of 0.15 mL/min. The column was held at 100% A for 2 minutes, ramped to 64% B over 18 minutes, ramped to 100% B over 1 minute, held at 100% B for 5 minutes, and equilibrated at 100% A for 25 minutes (50 minutes total). The column was maintained at 30 °C. The injection volume was 2 µL for samples and standard mixes. When starting a batch, the column was equilibrated at the starting conditions for at least 30 minutes. To improve the performance of the HILIC column, we maintained the same injection volume, kept the instrument running water blanks between samples as necessary, and injected standards in a representative matrix (the pooled sample) in addition to standards in water. After each batch, the column was flushed with 10 mM ammonium carbonate in 85:15 water to acetonitrile for 20 to 30 minutes.

The Waters Acquity UPLC was coupled to a Thermo Q Exactive HF hybrid Orbitrap high resolution mass spectrometer equipped with a heated electrospray ionization source (H- ESI). The H-ESI voltage was set to 3.3 kV and sheath gas, auxiliary gas, and sweep gas flow rates were set at 16, 3, and 1, respectively. The capillary and auxiliary gas heater temperatures were maintained at 320°C and 100°C, respectively. Full scan analyses were performed with polarity switching and a scan range of 60 to 900 *m/z* at a resolution of 60,000. The instrument was mass calibrated at 200 *m/z* before each run began and once again during the eddy transect sample set to ensure calibrations were always performed within 3-4 days. All data files were then converted to an open-source mzML format and vendor centroided via Proteowizard’s msConvert tool. All mzML files have been uploaded to Metabolomics Workbench under Project ID PR001738.

### Targeted metabolomic methods

Compounds for which we had an authentic standard (Supplemental Table 2) were manually integrated using Skyline and MSDIAL with the standard mixes used to ensure the correct peak was integrated by matching retention time. Compounds that were visually determined to have poor peak quality in the samples were removed from the downstream analysis. Skyline was used for both the eddy center samples and the HEE cruise data while MSDIAL was used for the eddy transect data due to the larger number of files in that dataset. Raw peak areas were then normalized to their best-matched internal standard (Supplemental Table 1) per Boysen et al. (2018) except that all compounds were normalized to their best match no matter how minimal the improvement. Normalized peaks were then calibrated via a single-point standard curve as determined uniquely for each compound from the authentic standard mixes and used to convert normalized peak area into moles per µL injected and scaled to the amount of seawater filtered to provide a final estimate of environmental concentration for each compound in moles per liter of seawater filtered.

### Untargeted metabolomic methods

Samples were also processed via the untargeted workflow detailed in Kumler et al. (2023). In short, XCMS (Smith et al. 2023) was used for feature detection with CentWave peakpicking, Obiwarp alignment, and peak density correspondence/grouping. This was followed by manual inspection of the extracted ion chromatogram for each mass feature by an expert for quality with only those peaks classified as “Good” used in the analysis here.

Features were then matched with and normalized to an internal standard (Supplemental Table 1) as described above.

### Statistics

All statistics were run in R version 4.3.3. We used multivariate statistics provided by the vegan package (Oksanen et al. 2022) to assess the overall impact of various metrics across the metabolome followed by univariate statistics to investigate individually significant responses. Non-parametric tests were used when data were unlikely to obey parametric assumptions and permutational statistics were preferred wherever possible. Multiple comparison problems were controlled where necessary using a false-discovery rate correction (Benjamini and Hochberg 1995). For the analyses involving the eddy transect samples, sea level anomaly was treated as a continuous variable with the exception of the PERMANOVA, for which SLA was categorized as cyclonic if less than -5 cm and anticyclonic if greater than 5 cm, with a third category for stations in between (Anderson 2001). Both the MESO-SCOPE eddy center and HEE analyses treated SLA as a categorical variable.

## Supplement

### Supplemental methods: Trichodesmium bloom in MS11C215mA?

One sample, taken from 15 meters at Station 11, was highly distinct from all others (including its paired triplicates) with several metabolites measuring two or more orders of magnitude larger in peak area and concentration both before and after normalization to internal standards (Supplemental Figure 3). Heuristic exploration of potentially related masses identified a total of 14 compounds highly abundant in this one sample alone. The only identified compounds with this trend were taurine and acetyltaurine, though three others with *m/z* ratios of 146.1176, 148.0968, and 206.1388 were discovered to be likely isomers of known compounds (acetylcholine, hydroxyisoleucine, and dexpanthenol, respectively). One feature (*m/z* = 184.0637, RT = 7.0 minutes) was identified as choline sulfate using a second run of standards, while an additional mass feature at 168.0689 and eluting at 8.5 minutes was putatively flagged as taurine betaine. This mass was also one of the most highly enriched in the sample after taurine itself, with a peak area nearly 30 times that of the next largest sample. No clear explanation emerged for why some compounds were hyperabundant in this surface sample, though one possibility was the *Trichodesmium* bloom that was also sampled at this time (Dugenne et al. 2023).

We explored the possibility that a nearby *Trichodesmium* bloom was responsible for the unusual composition of this one sample by analyzing additional samples taken directly from the bloom to see whether compounds abundant in hand-picked *Trichodesmium* colonies were also abundant in the Station 11 sample. After searching for the 14 mass features above in the environmental sample, 7 had good-quality peaks when viewed in the bloom samples as extracted ion chromatograms, though only 4 of these matched closely in retention time. We also performed a search of the untargeted data for homoserine betaine, a known *T. erythraeum* (strain IMS101) osmolyte (Pade et al. 2016). Although this compound was not run in water alongside the sample to confirm a retention time match due to a lack of authentic standards, we identified three mass features in the 162.1125 *m/z* window with retention times of 7.1, 8.4, and 9.9 minutes. The feature at RT 9.9 matched our carnitine standard’s retention time, leaving the two earlier peaks with putative identifications of threonine betaine at 7.1 and homoserine betaine at 8.4 given the elution order of threonine before homoserine in our authentic standards. This order is the reverse of that in Pade et al. (2016), which showed homoserine betaine eluting prior to carnitine. This elution order is expected given our use of a normal phase HILIC column and their use of a reversed phase Hypercarb column. They did not report an additional peak detected at this mass, leaving our putative identification of the homoserine and threonine betaines as low-confidence. None of the three 162.1125 features showed the same degree of enrichment in the anomalous sample as taurine or the other 13 compounds listed in Table 1, though the anomalous sample contained the largest, second-largest, and third-largest peak for carnitine, threonine betaine, and homoserine betaine, respectively. These results are fairly inconclusive but we are inclined to believe that a *Trichodesmium* colony is not likely to be the cause of the inflated values in this particular sample.

**Table 1:** Detected mass features that were enriched in the 15 meter surface sample from MESO-SCOPE Station 11. RT = retention time in minutes. Taurine betaine is followed by ato denote that this identification is putative. Mass difference between the median m/z and putative formula is reported in parts-per-million.

| Median<br>m/z | Median RT (min) | Putative formula | Delta mz (ppm) | Compound name |
| --- | --- | --- | --- | --- |
| 126.0221 | 11.1 | C2H8NO3S | -3.35 | Taurine |
| 146.1176 | 8.5 | C7H16NO2 | -3.63 |  |
| 148.0968 | 8.5 | C6H14NO3 | -3.88 |  |
| 168.0317 | 4.9 | C4H10NO4S | -7.87 | N-Acetyltaurine |
| 168.0689 | 8.5 | C5H14NO3S | -3.33 | Taurine betaine? |
| 176.1282 | 8.1 | C8H18NO3 | -2.88 |  |
| 184.0637 | 7.0 | C5H14NO4S | 0.00 | Choline sulfate |
| 192.1230 | 9.2 | C8H17NO4 | 0.00 |  |
| 196.0816 | 5.2 | C6H13NO6 | 0.00 |  |
| 198.0794 | 5.0 | C6H16NO4S | 0.00 |  |
| 206.1388 | 7.1 | C9H20NO4 | -1.93 |  |
| 208.1180 | 10.2 | C8H17NO5 | 0.00 |  |
| 222.1336 | 8.5 | C9H19NO5 | 0.00 |  |
| 266.1598 | 8.5 | C11H23NO6 | 0.00 |  |
| 266.1598 | 10.0 | C11H23NO6 | 0.00 |  |

---

**Supp fig 1:**
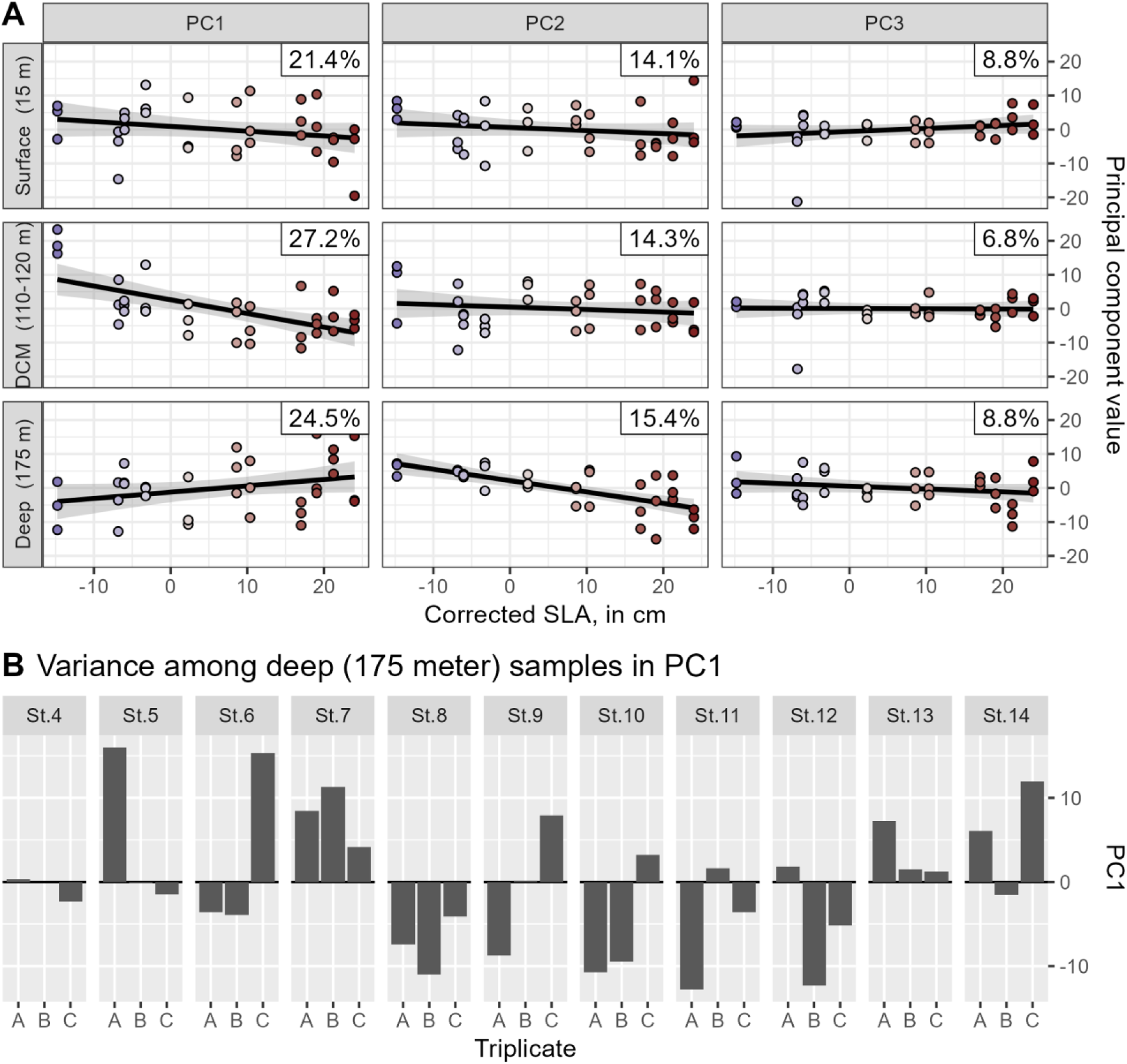
PCA axes 1-3 vs SLA by depth. Results of a principal component analysis performed separately on each depth of the eddy transect metabolome. A) Regressions of SLA against various principal components, broken down by depth. % variance explained by each PC is noted in the upper right corner of the plot. Colors have been added corresponding to the SLA value for clarity but are redundant with the x-axis values. Lines of best fit and the associated standard error have been added behind the points in black and translucent grey, respectively. Strong associations exist between PC1 and SLA in the DCM samples and PC2 and SLA in the 175 meter samples. B) Principal component 1 values for the 175 meter samples at each station showing large variance between triplicates rather than any external factor.

**Supp fig 2:**
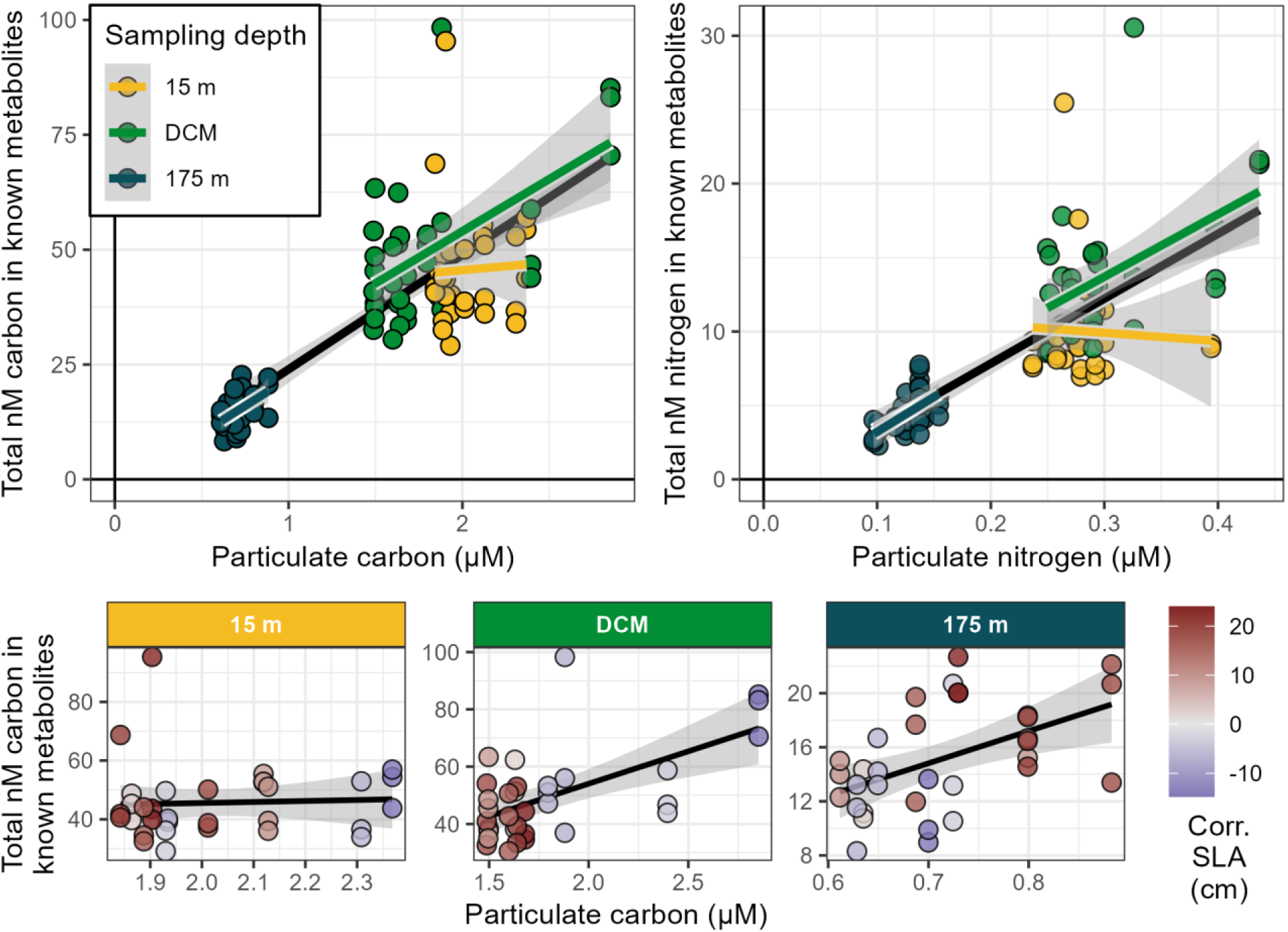
Biomass regression. Type I linear regressions of particulate carbon and nitrogen against total nM carbon and nitrogen in known metabolites both as a whole and divided by depth. Points have been colored according to depth (top row of plots) or SLA (bottom row of plots) and have been placed on independent axes. Two outlier particulate carbon points have been interpolated from beam attenuation for 15 meter values at stations 10 and 11.

**Supp fig 3:**
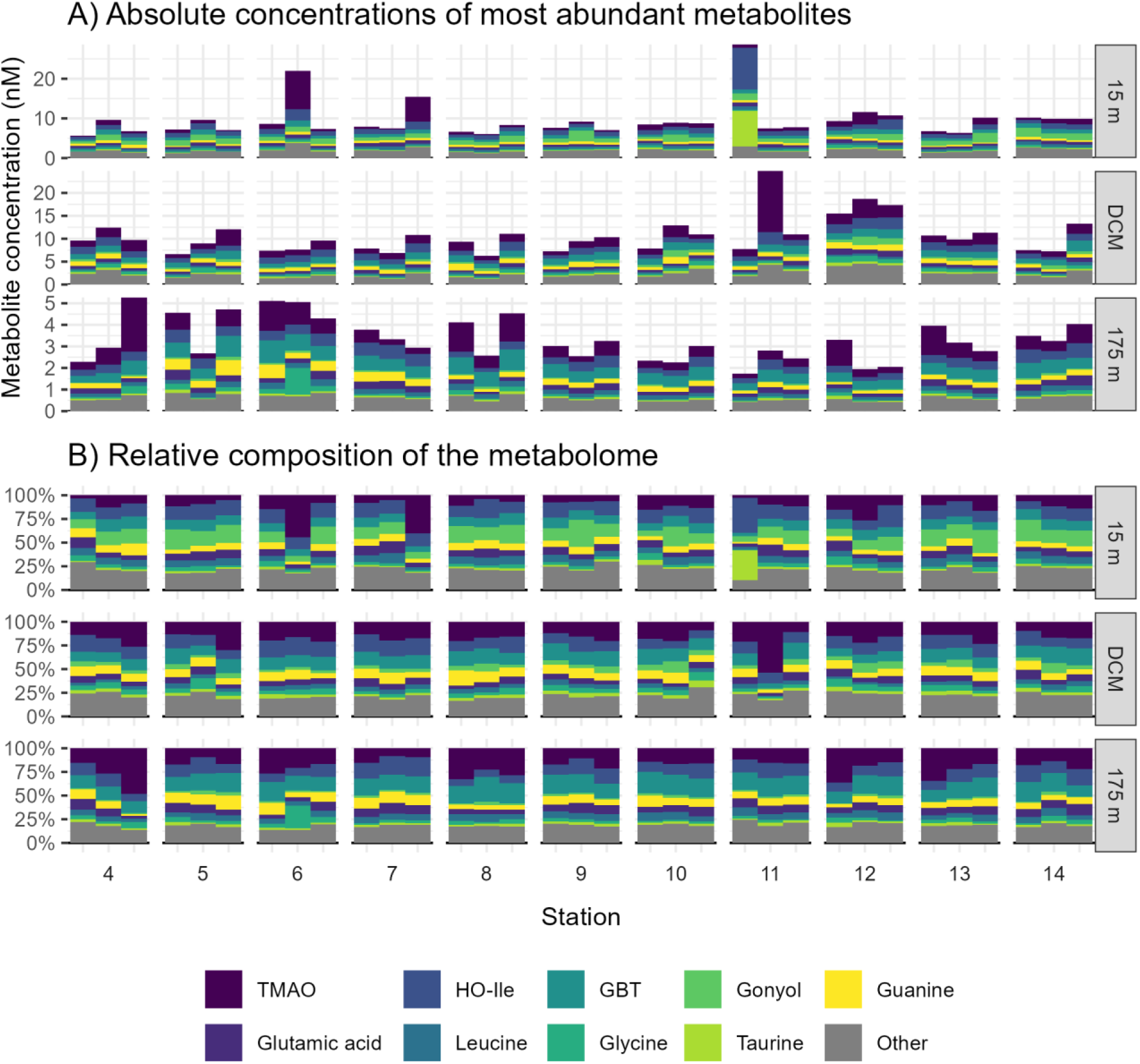
Stacked barplot of top metabs by triplicate. Stacked barplots of known metabolite concentration as shown in main text Figure 3 but with individual triplicates shown. The same data is shown in A) as B) but B) is rendered in relative contribution space instead of absolute concentration. TMAO = trimethylamine N-oxide, HO-Ile = hydroxyisoleucine, GBT = glycine betaine, DCM = deep chlorophyll maximum.

**Supp fig 4:**
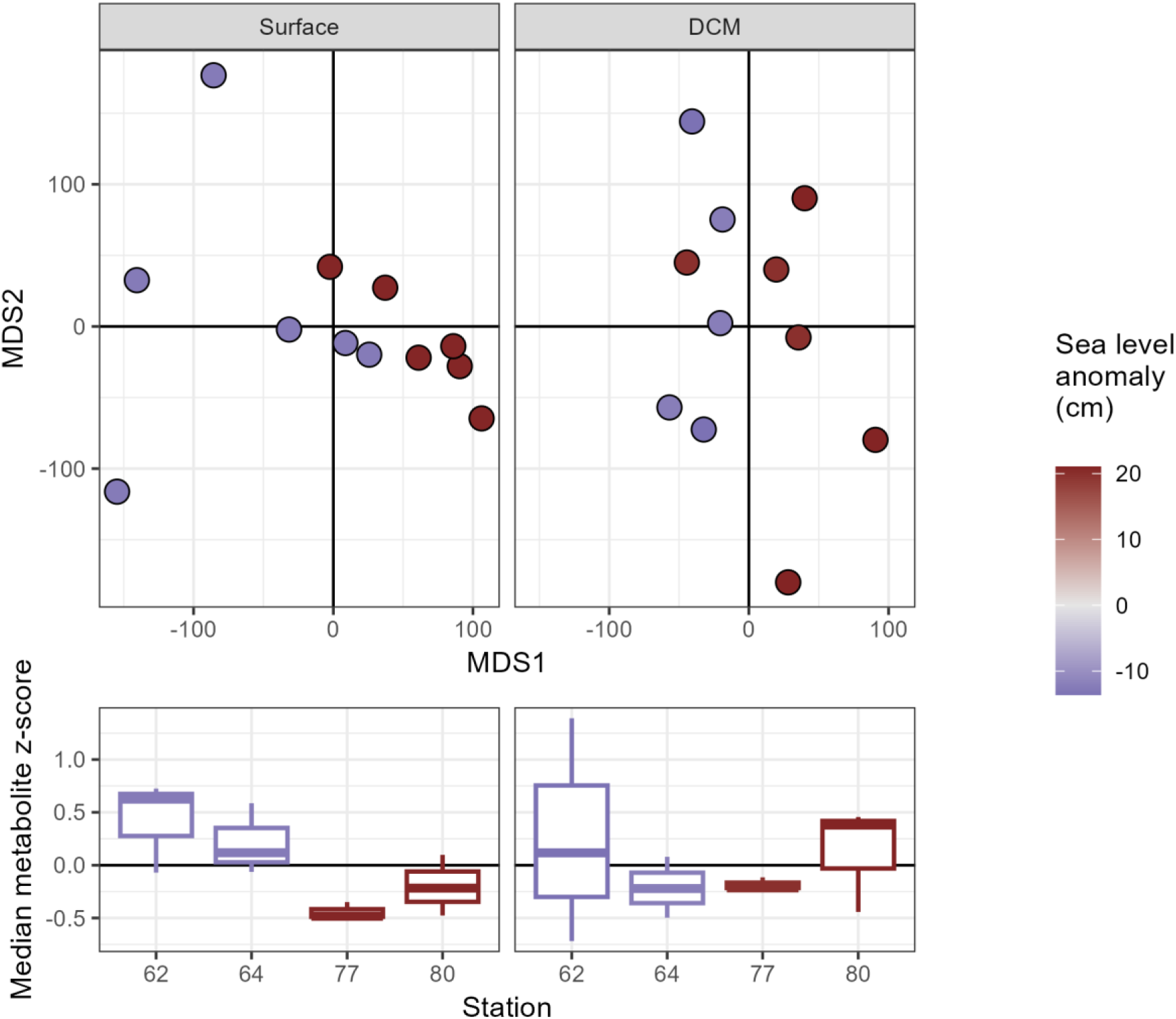
HEE NMDS and PERMANOVA broken down by depth. Plots of the metabolome during the Hawaiian Eddy Experiment, broken down by depth. The top row of plots are non-metric multidimensional scaling (NMDS) plots, in which points correspond to individual samples and have been colored by their corrected sea level height anomaly. SLA trends are visible in the 25 meter samples, with dark blue circles consistently discriminating from the dark red circles. The bottom row of plots show the direction and magnitude of this effect by plotting the grand mean of the normalized metabolite peak areas in the stations taken between the adjacent eddies of opposite polarity.

